# *AhRLK1*, a *CLAVATA1*-like leucine-rich repeat receptor-like kinase of peanut, confers increased resistance to bacterial wilt in tobacco

**DOI:** 10.1101/430066

**Authors:** Chong Zhang, Hua Chen, Rui-Rong Zhuang, Yuting Chen, Ye Deng, Tie-Cheng Cai, Shuai-Yin Wang, Qin-Zheng Liu, Rong-Hua Tang, Shi-Hua Shan, Rong-Long Pan, Li-Song Chen, Wei-Jian Zhuang

## Abstract

Bacterial wilt caused by *Ralstonia solanacearum* is a devastating disease that infects hundreds of plant species. Host factors involved in disease resistance and pathogenesis remain poorly characterized. An up regulated and leucine-rich repeat receptor-like kinase characterized as *CLAVATA1* and named *AhRLK1* was obtained by microarray analysis in response to *R. solanacearum* in peanut. AhRLK1 contained presumably, a signal peptide, ten leucine-rich repeat (LRR) domains and conserved motifs of intracellular kinases. For subcellular localization, the AhRLK1 protein was visualized only in the plasma membrane. After inoculation with *R. solanacearum*, AhRLK1 was constantly up regulated in the susceptible variety Xinhuixiaoli but showed little changed in the resistant cultivar Yueyou92. Different hormones, including salicylic acid, abscisic acid, methyl jasmonate, and ethephon, induced expression, but expression was completely down regulated under cold and drought treatments. Transient overexpression provoked a hypersensitive response (HR) in *Nicotiana benthamiana* following agro-infiltration. Furthermore, in transgenic tobacco with overexpression of the gene, the resistance to *R. solanacearum* increased significantly. By contrast, most representative defense-responsive genes in HR, SA, JA and ET signal pathways such as *NtHIN1, NtPR2, NtLOX1*, and *NtACS6*, among others, were considerably up regulated in the *AhRLK1* transgenic lines. Additionally, the *EDS1* and *PAD4* in the R gene signal were also up regulated in transgenic plants, but the *NDR1* and *NPR1* genes were down regulated. Accordingly, we suggest that *AhRLK1* increases the defense response to *R. solanacearum* via HR and hormone defense signalling, associated with the *EDS1* pathway of R gene signalling. The results provide new understanding of *CLV1* function and will contribute to genetic enhancement of peanut.

## Introduction

Bacterial wilt (BW) caused by *Ralstonia solanacearum* is a severe plant disease worldwide. The hosts of *R. solanacearum* include over 460 species in 54 botanical families (Wicker *et al*., 2007). As a soilborne disease, BW reduces peanut output in infected areas by 10–30 %, thereby causing significant economic loss and even leading to total crop failure in heavily infected regions. To date, efficient methods to control BW remain unavailable for all plants. Rotation, intercropping with other non-host crops, and biological control can help prevent BW incidence temporarily. However, the breeding of cultivars with genes resistant to BW infection is highly preferred.

Plants evolved a multi-layered innate immune system to defend against pathogens. Pattern recognition receptors (PRRs) on the plant cell surface act as initial detectors recognizing pathogen-associated or damage-associated molecular patterns to elicit the first-layer immune response called PAMP-triggered immunity (PTI) (Jones and Dangl, 2006; Zipfel, 2014). Thus, PTI prevents infections of non-adapted pathogens. Some adapted and successful pathogens deploy effectors that contribute to their virulence. Effectors subsequently interfere with PTI and cause effector-triggered susceptibility. In resistant plants, these effectors are recognized by R proteins to induce effector-triggered immunity (ETI) (Jones and Dangl, 2006). The co-evolution of PTI and ETI has reciprocally shaped the plant immune system (Böhm *et al*., 2014).

Most PRRs are characterized as leucine-rich repeat receptor-like protein kinases (LRR-RLKs) that compose a class of RLKs in plants (Zhang, 1998). Presumably, LRR-RLK-encoded proteins participate in the regulation of plant growth and development (Morris and Walker, 2003), hormone signal transduction (Hong *et al*., 1997), and biotic or abiotic stress responses (Nishiguchi *et al*., 2002; Torii, 2004). LRR-RLKs are also involved in plant defense-related disease resistance (Song *et al*., 1995; Godiard, Laurence and Sauviac, Laurent and Torii, Keiko U and Grenon, Olivier and Mangin, Brigitte and Grimsley, Nigel H and Marco, 2003). A typical LRR-RLK structure is composed of extracellular domains (LRR), single transmembrane domains flanked by juxta membrane regions, and cytoplasmic protein kinase domains (Dardick *et al*., 2012; Zhang and Thomma, 2013; Böhm *et al*., 2014). LRR domains function as binding sites for the specific recognition of pathogen-derived elicitors to activate downstream signal transduction by the cytoplasmic protein kinase domains, thereby enabling the plant to produce a defensive immune response (Jones and Jones, 1997; Dardick *et al*., 2012; Böhm *et al*., 2014).

FLAGELLIN SENSITIVE2 (FLS2), an LRR-RLK protein, is a plasma membrane receptor involved in the recognition of pathogen flagellin (Gómez-Gómez and Boller 2000). FLS2 has receptor activity (flagellin binding) in its extracellular domain, and the kinase domain is required to induce the pathogen response (Gómez-Gómez and Boller, 2000; Gómez-Gómez *et al*., 2001). Another LRR-RLK gene named *Xa21* is a resistance gene to leaf blight in rice (Wang *et al*., 1996). Xa21 is composed of 21 LRR motifs that recognize pathogen ligands, eliciting plant defense responses, such as oxidative bursts, hypersensitive cell death, and defense gene activation, via intracellular kinases (Song *et al*., 1995; Wang *et al*., 1996). *ERECTA* is another *Arabidopsis* LRR-RLK gene resistant to *R. solanacearum* (Godiard, Laurence and Sauviac, Laurent and Torii, Keiko U and Grenon, Olivier and Mangin, Brigitte and Grimsley, Nigel H and Marco, 2003), which activates the expression of downstream resistance-related genes against *R. solanacearum* infection by extracellular kinase phosphorylation (Godiard, Laurence and Sauviac, Laurent and Torii, Keiko U and Grenon, Olivier and Mangin, Brigitte and Grimsley, Nigel H and Marco, 2003). Additionally, *ERECTA* triggers a resistance response to necrotize fungi (*Plectosphaerella cucumerina*) in *Arabidopsis* (Llorente *et al*., 2005). An increasing number of LRR-RLKs are expected to be identified and their resistance mechanisms in plant–pathogen interactions to be explicitly elucidated.

In the present study, an LRR-RLK gene named *AhRLK1* was obtained from peanut by microarray analysis. The *AhRLK1*, characterized as *CLAVATA 1*, was up regulated in a peanut cultivar susceptible to BW but remained nearly unchanged in the resistant one. Different hormones and cold or drought treatments induced the expression of this gene. Transient overexpression caused a hypersensitive response (HR) in *Nicotiana benthamiana* following agro-infiltration. Furthermore, with overexpression of *AhRLK1* in *Nicotiana tabacum*, resistance to *R. solanacearum* increased significantly. The expression levels of various stress-responsive genes including those of R gene signalling were also significantly up regulated in the *AhRLK1*-overexpressing transgenic lines. Therefore, these results suggest that *AhRLK1* is involved in the defense response of peanut to *R. solanacearum* and in the resistance conferred by multiple, complex signalling regulatory networks.

## Materials and methods

### Plant materials and growth conditions

The Oil Crop Institute of Fujian Agriculture and Forestry University provided the peanut (*Arachis hypogaea*) cultivars that were middle resistant (Minhua 6), hyperresistant (Yueyou 92), and hypersusceptible (Xinhuixiaoli) to *R. solanacearum*. Seeds were sown in sterile sand in 5 × 6 cm plastic pots. The Tobacco Research Group of Fujian Agriculture and Forestry University provided the seedlings of transgenic and wild-type tobacco lines (*Nicotiana tabacum* cv. CB-1, cv. Honghuadajinyuan, and cv. Yanyan97 with medium susceptibility, hypersusceptibility, and hyperresistance to *R. solanacearum*, respectively) and those of *N. benthamiana*. All seedlings were grown in a greenhouse. T_1_ and T_2_ seeds of transgenic tobacco lines were surface-sterilized with 75 % (w/v) alcohol for 20 s and 10 % (v/v) H_2_O_2_ for 10 min, washed five times with sterile water, and then placed on MS medium supplemented with 75 mg/L kanamycin for 2–3 weeks. The surviving plants were transferred into a soil mix (peat moss/perlite, 2/1, v/v) in a plastic tray and grown in a greenhouse for another 2–3 weeks. Transgenic and wild-type tobacco plants of the same size were transferred into the same soil mixed in plastic pots and grown for another 3–4 weeks. The peanut and tobacco plants were grown in a greenhouse at 26 ± 2 °C, with 70 % relative humidity and a 16 h-light/8 h-dark cycle.

### Pathogens and inoculation procedures

Virulent *R. solanacearum* strains were used in this study, i.e., *Rs-P.362200–060707-2–2* for peanut and FJ1003 for tobacco. The pathogen strain was streaked on TTC agar medium (0.5 g/L 2,3,5-triphenyltetrazolium chloride, 5 g/L peptone, 0.1 g/L casein hydrolysate, 2 g/L D-glucose, and 15 g/L agar) (Kelman *et al*. 1954) and then incubated at 28 °C for 48 h. Virulent colonies, white clones with pink centers, were harvested with sterile water containing 0.02% Tween-20, and the inoculum was prepared by adjusting the concentration of bacterial cells to an optical density of 0.5 at 600 nm (Nano Drop 2000c; Thermo Fisher Scientific, Middletown, VA, USA). This optical density corresponded to approximately 10^8^ colony-forming units (cfu)/mL for inoculating peanut and tobacco seedlings. After 4 weeks, the third and fourth leaves from the upper part of peanut seedlings of Yueyou92 and Xinhuixiaoli were inoculated by leaf cutting per leaflet (perpendicularly to the midrib, up to a 2/3 portion), with four leaflets per branch. Control plants were inoculated with distilled water containing 0.02 % Tween-20. Two uncut leaflets per leaf were harvested at the indicated time points and used as an RNA source for future analysis. For tobacco inoculation, 10 µL of the *R. solanacearum* suspension (10^8^ cfu/mL) was infiltrated into the third leaf from the top using a syringe with a needle, and then the fourth leaf was harvested at the indicated time points and used as an RNA source for future analysis. Typical symptoms of BW were monitored daily with five disease severity scores that ranged from 0 to 4, where 0 = no symptoms, 1 = 1/4 of inoculated leaves wilted, 2 = 1/4−1/2 of inoculated leaves wilted, 3 = 1/2−3/4 of inoculated leaves wilted, and 4 = whole plant wilted, with plant death. Disease index (DI) and death ratio (DR) were calculated using the following formulas: DI (%) = [∑ ni × vi) ÷ (*V* × *N*)] × 100 and DR (%) = (ni ÷ *N*) × 100, where ni = number of plants with the respective disease rating; vi = the disease rating; V = the highest disease rating; and N = the total number of observed plants.

For the transient overexpression of *AhRLK1* in *N. benthamiana*, 10^8^ cfu/mL *Agrobacterium* was infiltrated into the second leaf of two-month-old tobacco from the top using a syringe without a needle until the bacterial suspensions had spread over the entire leaf. The third leaf was harvested at the indicated time points, immediately frozen in liquid nitrogen, and then stored at −80 °C for further use.

### Application of plant hormones or abiotic and biotic stresses

One-month-old peanut (Minhua 6) seedlings were sprayed with 3 mM salicylic acid (SA), 10 µg/mL abscisic acid (ABA), 10 mM ethephon (ET), or 100 µM methyl jasmonate (JA) in distilled water. Control seedlings were sprayed only with distilled water. At various time intervals, the leaves of the treated seedlings were harvested, frozen in liquid nitrogen, and then stored at −80 °C until further use. Peanut (Minhua 6) plants at the seven-leaf stage were treated at a low temperature of 4 °C or a normal temperature of 25 °C. Leaves were harvested at the indicated time points. For drought stress, peanut (Minhua 6) plants at the seven-leaf stage were treated either without watering or with normal watering. Leaves were harvested at different time intervals. All samples had three biological replicates; these samples were frozen in liquid nitrogen and then stored at −80 °C until further use.

### Full-length cDNA cloning of AhRLK1

As a candidate differentially expressed gene, the *AhRLK1* fragment was screened using a high-density peanut microarray with a hundred thousand unigenes, which was devised by our laboratory and created by the Roche Company (Roche, Branford, Connecticut, USA). The *AhRLK1* gene was isolated by chip hybridization using RNAs extracted from peanut plants with/without inoculation of *R. solanacearum*. For cloning of full-length *AhRLK1*, AhRLK1-F and AhRLK1-R primers were designed from the available gene fragments. The 5′- and 3′-end sequences of the cDNA were cloned through RACE using a SMART^™^ RACE cloning kit (Clontech, Palo Alto, CA) according to the manufacturer’s instructions with minor modifications. Total RNA was extracted from the leaves of the peanut cultivar hyperresistant to *R. solanacearum* using the CTAB method (Chen *et al*., 2016). The adaptor primers of RACE-F and 3′ PCR primer were ligated to both ends of the cDNA. The 5′ RACE was generated by PCR with the primary primer set of RACE-F primer and AhRLK1-R. The reaction condition was as follows: 94 °C for 5 min; 35 cycles of 95 °C for 30 s, 60 °C for 30 s, and 72 °C for 1.5 min; and 72 °C for 10 min. Similarly, the 3 ′ RACE was generated by the set of AhRLK1-F and the 3′ PCR primers. The PCR program was as follows: 94 °C for 5 min; 5 cycles of 95 °C for 30 s and 72 °C for 2 min; 30 cycles of 95 °C for 30 s, 60 °C for 30 s, and 72 °C for 2 min; and 72 °C for 10 min. The RACE products were ligated to pMD18-T vectors (TaKaRa Biotech. Co., Dalian, China) in accordance with the manufacturer’s instructions and then sequenced. After assembly, the full-length cDNA sequence and DNA sequence of *AhRLK1* were cloned from the reverse transcription products and genomic DNA by using AhRLK1-FL-F and AhRLK1-FL-R. All the primers employed in this study are listed in supplemental table S1.

### Sequence analysis and phylogenetic tree construction

*AhRLK1* sequence similarity analysis was performed with BLASTN and BLASTX (http://www.ncbi.nlm.nih.gov/BLAST). Conserved domains of the *AhRLK1-* encoded protein were analysed using SMART (Simple Modular Architecture Research Tool) (http://smart.embl-heidelberg.de/). Multiple sequence alignments were obtained from known functional LRR–RLKs of different species using Clustal 2W. A phylogenetic tree involving different subfamilies of LRR-RLKs in *Arabidopsis* was generated with the MEGA 5.10 program (Tamura *et al*., 2011).

### Subcellular localization

The full-length open-reading frame of *AhRLK1* without the termination codon was amplified by high-fidelity PCR polymerase with pMD-T-AhRLK1 as the template. The gene-specific primers AhRLK1-BamH1-F and AhRLK1-Asc1-R harbouring *Bam* HI and *Asc*I sites, respectively, were employed. The PCR products and the pBI-green fluorescent protein (GFP) vector were both digested with *Bam* HI and *Asc*I. The corresponding bands were recovered and ligated to the 35S::AhRLK1-GFP expression vector. The 35S::GFP vector was used as a control and transformed into *Agrobacterium* strain GV3101. The *Agrobacterium* strain GV3101 harbouring the above mentioned constructs was grown for 24 h in YEP medium (10 g/L yeast extract, 10 g/L peptone, and 5 g/L NaCl) containing appropriate antibiotics. *Agrobacterium* was suspended in infiltration buffer (10 mM MgCl_2_, 10 mM 2-(*N*-morpholino) ethanesulfonic acid, and 200 mM acetosyringone, pH 5.7). *Nicotiana benthamiana* leaves were infiltrated with the infiltration cultures. After 2 days of infection, GFP fluorescence was visualized under a fluorescence microscope at a 488 nm excitation wavelength and a 505–530 nm band pass emission filter. Digital images were overlaid using Image-J.

### AhRLK1 overexpression vector construction, transient expression, and tobacco transformation

The complete ORF of AhRLK1 was amplified by high-fidelity PCR polymerase with pMD-T-AhRLK1 as the template. The primers AhRLK1-OE-F and AhRLK1-OE-R harbouring *Bam* HI and *Asc* I sites, respectively, were employed. The PCR products and the pBI121-GUSA vector were both digested with *Bam* HI and *Asc* I; the corresponding bands were recovered and ligated into pBI121-GUSA driven by the 2×CaMV 35S promoter to generate the overexpression vector 35S::AhRLK1. The 35S::AhRLK1 plasmid was transferred into *Agrobacterium tumefaciens* strains GV3101 and EHA105. For transient expression, *Agrobacterium* GV3101 with the 35S::AhRLK1 plasmid was injected into *N. benthamiana* leaves via *Agrobacterium* infiltration and then transformed into tobacco via the leaf-disc method (Müller *et al*., 1987). To confirm transgene integration, the initial transgenic T_0_ lines were selected by kanamycin and further confirmed by reverse transcription-PCR (RT-PCR). The T_2_ pure lines were obtained and used in this study.

### In silico analysis and quantitative Real-Time PCR

In silico analysis of *AhRLK1* gene expression pattern in peanut was performed using non-amplified double strain cDNA for hybridization as described previously (Chen *et al*. 2016). The gene expression intensity of all hybridizations was analysed, and expression levels were estimated among different tissues and under diverse stress conditions. Three replicates were performed for all experiments.

The data from the tobacco microarray were determined previously (Zhang et al., 2017). Leaves were harvested of the hyperresistant tobacco variety Yanyan 97 and the hypersusceptible tobacco variety Honghuadajinyuan after *R. solanacearum* inoculation. Microarray design, hybridization, washing, and scanning and data analysis were conducted as previously described (Zhang et al., 2016).

For qRT-PCR analysis, total RNA was extracted from peanut, transgenic tobacco, and wild-type seedlings using the CTAB extraction method (Chen *et al*. 2016). A 1 µg RNA sample was reverse transcribed with PrimeScript^™^ RTase in accordance with the manufacturer’s instructions (TaKaRa Biotech. Co., Dalian, China). The cDNA was then diluted to 1:10 with diethylpyrocarbonate-treated H_2_O before use. Real-time PCR for the relative expression level of the target gene was performed with specific primers (Table 1 lists the gene-specific primers) that were essentially provided for a Mastercycler eprealplex (Eppendorf, Hamburg, Germany) and SYBR Premix Ex Taq II (perfect real time; TaKaRa Biotech. Co., Dalian, China). Each reaction mixture (20 µL) contained 10 µL of SYBR Premix Ex Taq (2×), 0.2 µL of PCR forward/reverse gene-specific primers (10 µM), and diluted cDNA (2 µL). For each gene, three experimental replicates were obtained with different cDNAs synthesized from three biological replicates. The PCR program was as follows: 95°C for 5 min; 40 cycles of 95°C for 5 s, 60°C for 30 s, and 72°C for 30 s; 95°C for 15 s, 60°C for 1 min, 95°C for 15 s, and 60°C for 15 s. The specificity of amplification was confirmed by melting curve analysis after 40 cycles. The relative expression level of the target gene was calculated via the comparative CT method (2^-ΔΔCT^ method) (Schmittgen and Livak, 2008) by normalizing the PCR threshold cycle number (Ct value) of the target gene with the reference gene. The calculation formula was ΔΔCt = (CT_gene_ – CT_actin_)_treat_ – (CT_gene_ – CT_actin_)_control_. The relative transcript levels of *AhRLK1* were detected under different treatments in peanut, with *Ahactin* as the internal reference. The relative transcript levels of related defense genes after *R. solanacearum* treatment were detected between the wild type and transgenic tobacco plants, with tobacco *NtEF1*α as the internal reference. All primers used for qPCR are listed in supplementary table S1.

**Table 1.** Comparison of disease index and death ratio of different OE lines and the wild type after inoculation with *R. solanacearum*

| OE Lines | 7 dpi* |  | 21 dpi |  |
| --- | --- | --- | --- | --- |
|  | Disease index | Death ratio | Disease index | Death ratio |
|  | (%) | (%) | (%) | (%) |
| Wild type | 72.97 | 27.91 | 95.64 | 86.05 |
| OE-1 | 14.32** | 0.00 | 21.84** | 6.80 |
| OE-7 | 28.79** | 6.98 | 42.73** | 12.79 |
| OE-19 | 34.76** | 7.32 | 58.84** | 45.12 |
| OE-32 | 26.09** | 9.78 | 34.51** | 11.96 |
| OE-43 | 27.60** | 7.79 | 41.56** | 23.38 |
| OE-46 | 31.25** | 11.96 | 54.62** | 30.43 |

\* dpi: days postinoculation; \*\* indicates highly significant difference.

**Table 1** Comparison of disease indices and death ratios of different OE lines and wild-type after inoculation with *R. solanacearum*
| OE Lines | 7 dpi* |  | 21 dpi |  |
| --- | --- | --- | --- | --- |
|  | Disease index | Death ratio | Disease index | Death ratio |
|  | (%) | (%) | (%) | (%) |
| Wild-type | 72.97 | 27.91 | 95.64 | 86.05 |
| OE-1 | 14.32** | 0.00 | 21.84** | 6.80 |
| OE-7 | 28.79** | 6.98 | 42.73** | 12.79 |
| OE-19 | 34.76** | 7.32 | 58.84** | 45.12 |
| OE-32 | 26.09** | 9.78 | 34.51** | 11.96 |
| OE-43 | 27.60** | 7.79 | 41.56** | 23.38 |
| OE-46 | 31.25** | 11.96 | 54.62** | 30.43 |

\* dpi: days post inoculation; \*\* represent extremely significant difference.

### Histochemical staining analysis and ion conductivity determination

At 48 h after the transient overexpression of AhRLK1 in *N. benthemiana* leaves, the infected plants were stained with 3,3′-diaminobenzidine (DAB; Sigma, St. Louis, MO) and lactophenol–ethanol trypan blue. To measure the levels of H_2_O_2_, the infected *N. benthemiana* leaves were incubated in 1 mg/mL DAB solution overnight at room temperature, boiled for 5 min in a 3:1:1 ethanol/lactic acid/glycerol solution, and then placed in absolute ethanol before observation. For cell death detection, the inoculated leaves were boiled in trypan blue solution (10 mL of lactic acid, 10 mL of glycerol, 10 g of phenol, 30 mL of absolute ethanol, and 10 mg of trypan blue, dissolved in 10 mL of ddH_2_O) for 2 min, left at room temperature overnight, transferred into a chloral hydrate solution (2.5 g of chloral hydrate dissolved in 1 mL of distilled water), and then boiled for 20 min for de-staining. The leaves were observed under a light microscope. Ion conductivity was measured as previously described with minor modifications (Hwang and Hwang, 2011). Six round leaf discs (11 mm in diameter) per agro-infiltrated leaf were cut, washed in ddH_2_O, and then incubated in 20 mL of ddH_2_O with evacuation for 10 min at room temperature. Electrolyte leakage was measured with a Mettler Toledo 326 apparatus.

## Results

### Sequence characteristics of AhRLK1 isolated from peanut

The 5′ and 3′ unknown cDNA sequences of *AhRLK1* were cloned by RACE. The full-length cDNA sequence was isolated from the total RNA of peanut leaf with RT-PCR (Fig. S1). The full-length cDNA contained a 3,292 bp ORF encoding a polypeptide of 992 amino acids with 122 and 251 bps for 5′ and 3′ untranslated terminal regions (UTR), respectively (Fig. 1; Data S1). Sequence analysis showed the deduced AhRLK1 protein contained the typical serine/threonine protein kinase catalytic domain and 10 LRR conserved domains (LxxLxxLxxLxLxxC/A-xx) (Leah McHale et al., 2012) (Fig. 1; Fig. S2). Additionally, the protein had a signal peptide in the N-terminal (Fig. 1).

**Figure 1.**
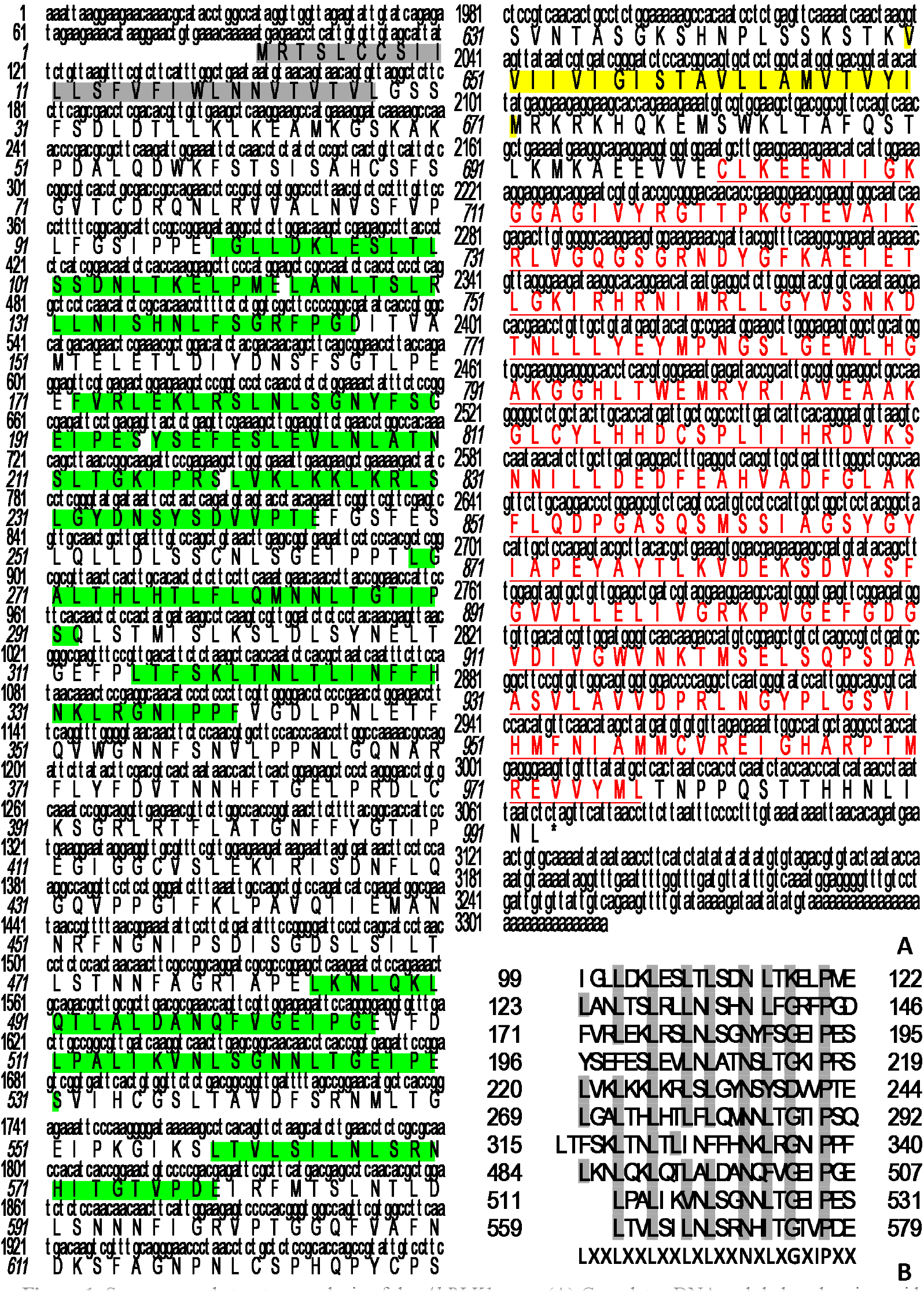
Sequence and structure analysis of the *AhRLK1* gene. (A) Complete cDNA and deduced amino acid sequences of the *AhRLK1* gene. The full-length cDNA was 3,292 bp with an ORF encoding 992 amino acids. The gray-shaded portion shows the signal peptide domain, and the green-shaded portion indicates the LRR units; the transmembrane domain is in the yellow-shaded region. The underlined red sequences show the serine/threonine protein kinase catalytic domain. (B) LRR domain of AhRLK1, including several degenerate LRR units. The consensus sequence for the AhRLK1 LRR is given at the bottom. The core leucines and prolines (or equivalent amino acids) are highlighted in gray. X represents an arbitrary amino acid residue. The L-residues in the consensus sequence represent several residues at that position.

A comparison study revealed that the amino acid sequence of AhRLK1 resembled that of resistance proteins of *Arabidopsis thaliana* CLAVATA1 (61 % identity and 75 % similarity), ERECTA (34 % identity and 50 % similarity) to *R. solanacearum*, BAM1 (50 % identity and 67 % similarity), and *A. thaliana* BAM2 (49 % identity and 67 % similarity) (Data S2; Fig. S2).

A phylogenetic tree for AhRLK1 further confirmed that AhRLK1 is a member of the LRR XI subfamily, with the most similarity to At1g75820 (Accession number: NP_177710), which encodes the CLAVATA1 (CLV1) protein (Fig. 2; Data S3; Fig. S3). CLV1 is involved in meristem differentiation and maintenance (Clark *et al*., 1997). However, these two kinases may have significantly diverged in their functions; thus, *AhRLK1* may be both a resistance and a structural gene.

**Figure 2.**
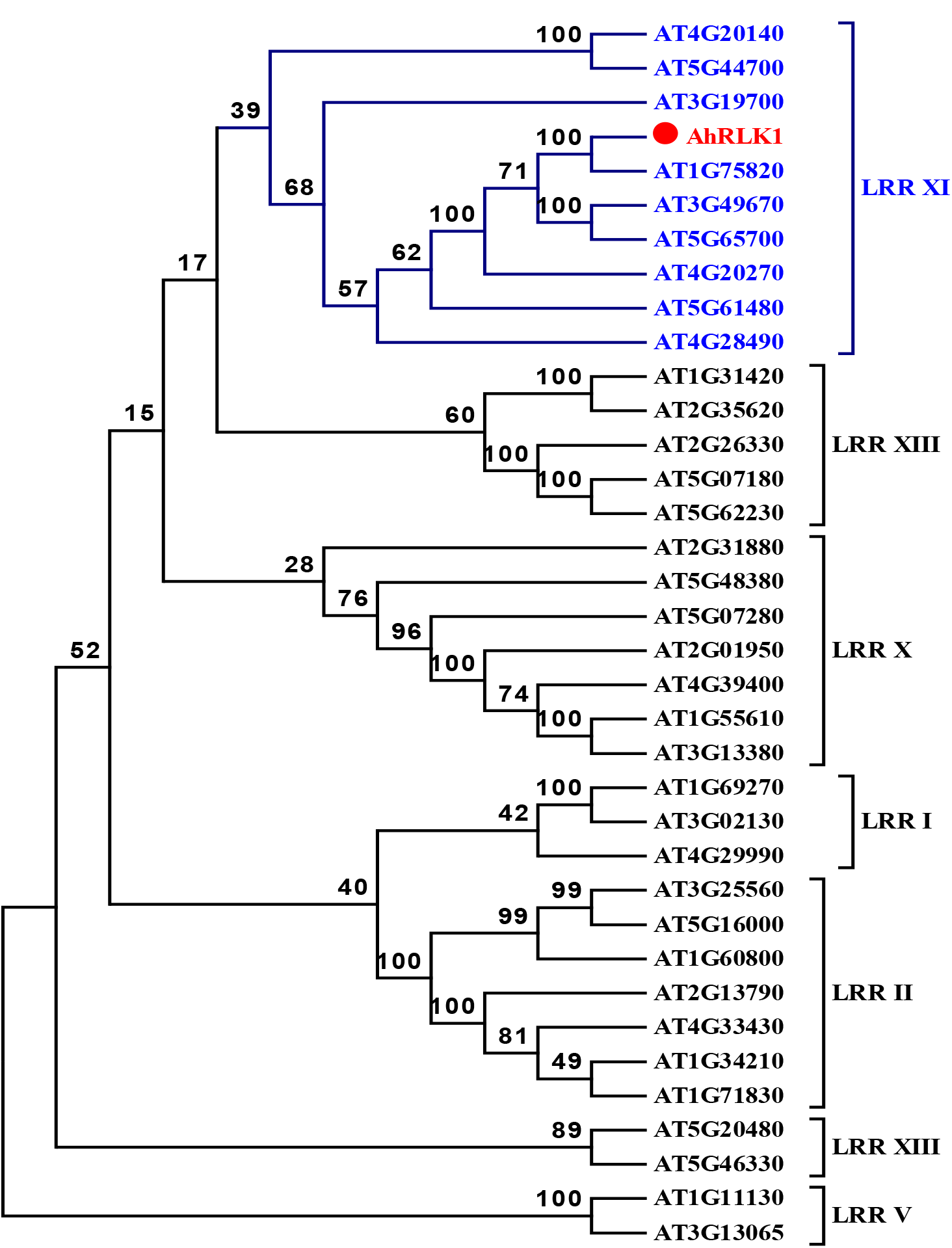
Phylogenetic tree constructed with AhRLK1 and different subfamilies of LRR-RLK proteins in *Arabidopsis*. The phylogenetic tree confirms that AhRLK1 is a member of the LRR XI family; AhRLK1 is indicated by a red rhombus. Alignments were conducted in ClustalW, and the phylogenetic tree was constructed by the neighbour-joining algorithm in MEGA 5.10 software. Bootstrap values (1,000 replicates) are shown as percentages at the branch nodes.

### Subcellular localization of AhRLK

Sequence analysis predicted that the AhRLK1 protein is a plasma membrane-bound kinase (Query Protein WoLFPSORT prediction plas: 29.29 by http://www.genscript.com/psort/wolf_psort.html). The subcellular localization of AhRLK1 was determined using a GFP fusion protein. The AhRLK1–GFP fusion protein driven by the constitutive CaMV35S promoter and a 35S::GFP as a negative control were generated. Both constructs were transformed into *Agrobacterium* strain GV3101, which was infiltrated into *N. benthamiana* leaves. Typical results indicated that the AhRLK1-GFP was localized in the plasma membrane and cytoplasm, whereas GFP alone occurred in multiple subcellular compartments, including the cytoplasm and nuclei (Fig. 3). Thus, AhRLK1 is a plasma and membrane-associated and cytoplasm kinase.

**Figure 3.**
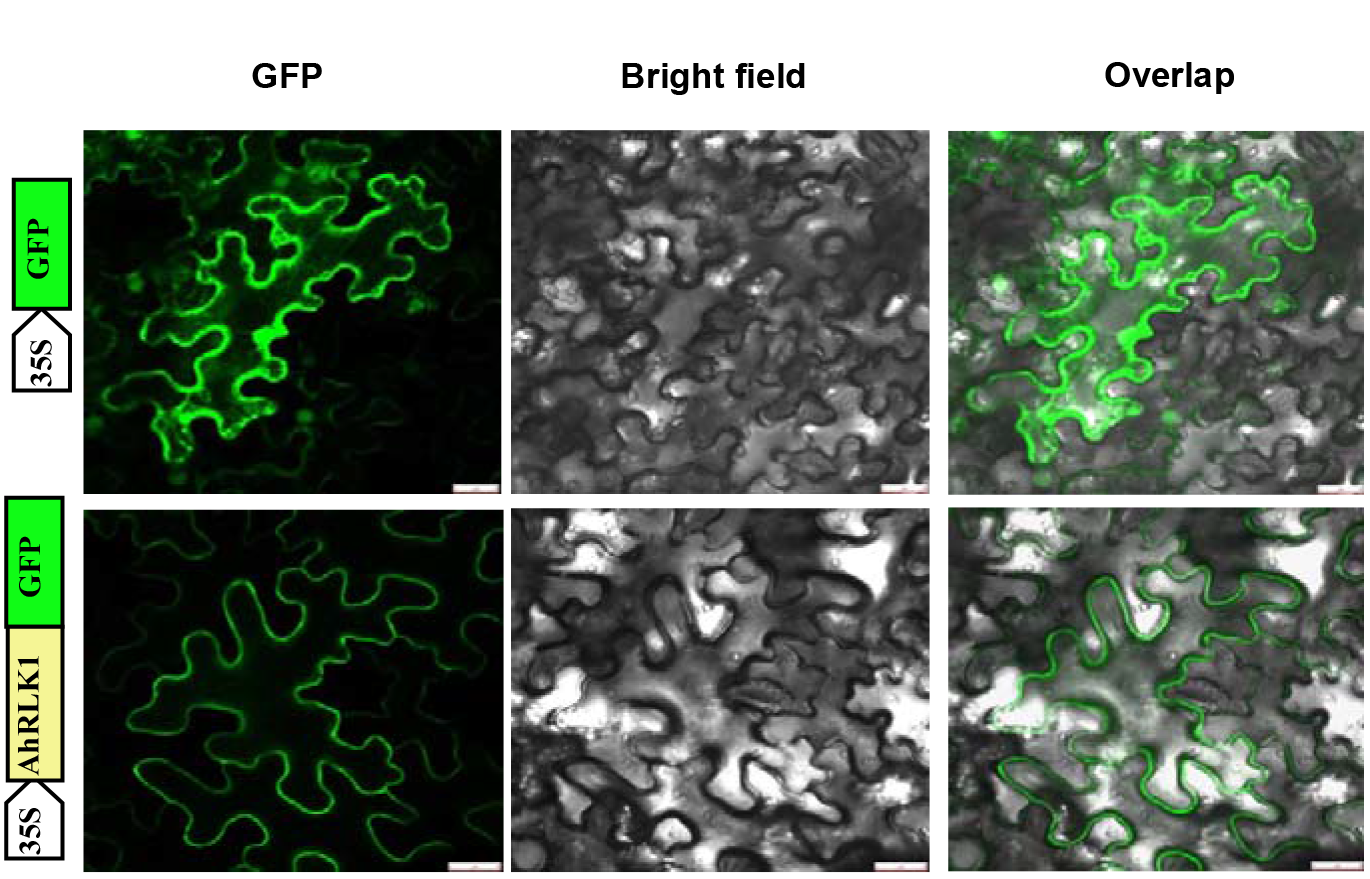
Subcellular localization of *AhRLK1*. AhRLK1-GFP was localized in the plasma membrane of *Nicotiana benthamiana* leaves; GFP alone was localized throughout entire cells. Fluorescence (left), bright field (middle), and merged images (right) were obtained at 48 h using Leica confocal microscopy after agro-infiltration. Bar = 25 μm.

### AhRLK1 showed diverse expression patterns among tissues

In silicon analysis of *AhRLK1* expression with two unigenes was performed using a high-density microarray. These unigenes with more than 97 % sequence identity apparently belonged to the same *AhRLK1* gene family. Non-amplified double strains of cDNA were employed to evaluate the transcript levels of the unigenes in the microarray hybridization. The expression profiles of these three members all demonstrated a harmonized pattern among tissues, with the highest expression in the roots and stem, followed by the leaves, flowers, pegs and testa. However, expression was weak in the pericarp, and embryos displayed the lowest expression levels of these genes (Fig. 4; Data S4).

**Figure 4.**
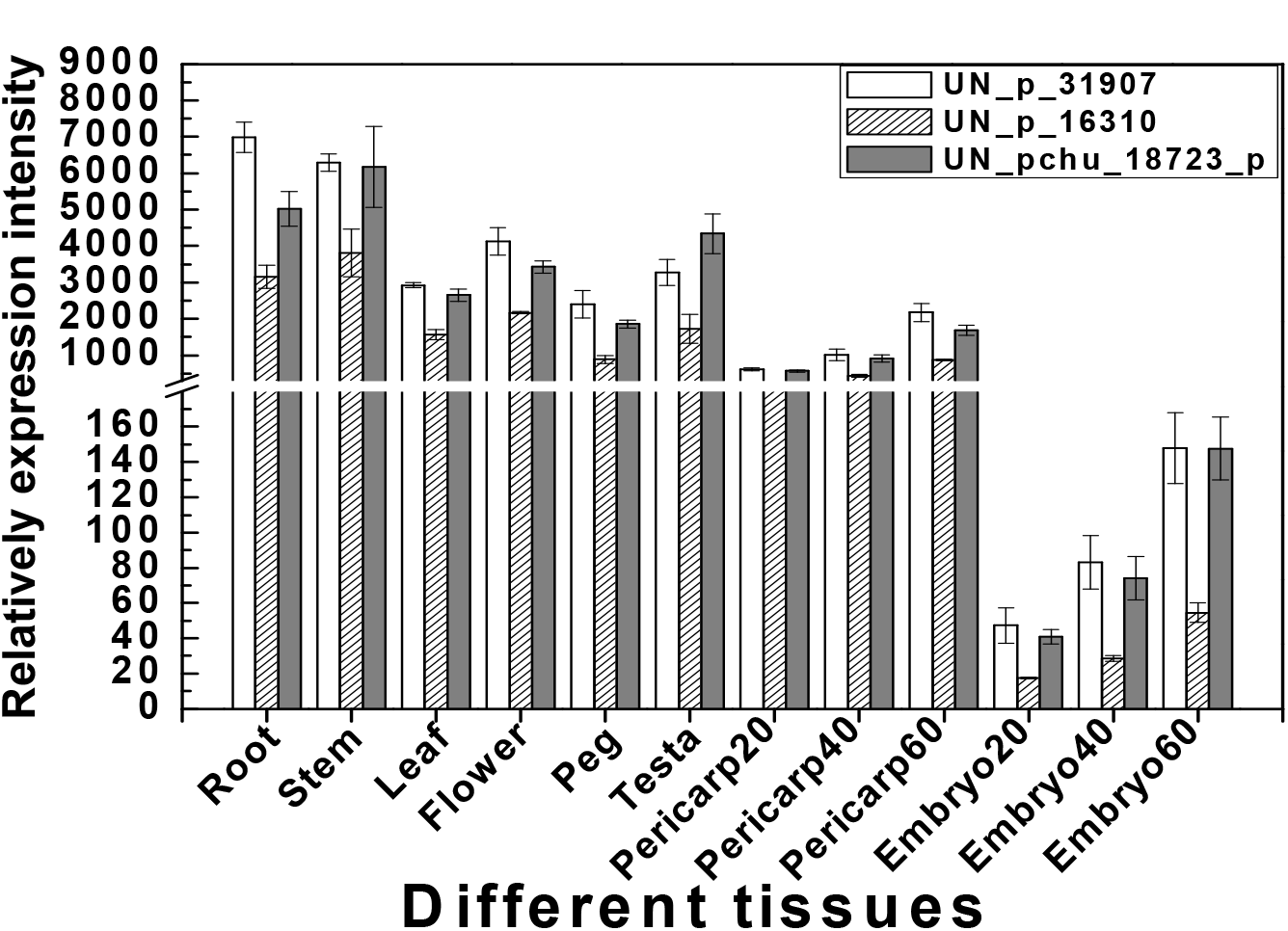
In silico identification and the expression characteristics of three members of the *AhRLK1* gene family. The *AhRLK1* family showed tissue-specific expression in peanut, with the highest levels in the roots and stem. Weak expression was found in pericarp and embryo. UN_p_31907, UN_p_16310 and UN_pchu_18723 are *AhRLK1* and the two other members of the same family, respectively.

### AhRLK1 responses to bio/abiotic stresses

The expression of *AhRLK1* under exogenous phytohormone treatments was determined in the medium resistance variety Minhua 6 at the eight-leaf stage (Fig. 5). Upon 3 mM SA treatment, the *AhRLK1* transcripts increased up to 6.6-fold at 6 hours post treatment (hpt) and then gradually decreased to levels slightly higher (<3 fold) than those of the control plants (Fig. 5A). The expression of *AhRLK1* also increased with 10 µg/mL ABA, reaching a single peak of 4.5-fold at 6 hpt (Fig. 5B). Similarly, after 100 µM MeJA treatment, *AhRLK1* expression also elevated progressively, with the highest level (3.8-fold up regulation) at 6 hpt (Fig. 5C). In response to 10 mM ET, *AhRLK1* expression increased with two peaks (2.6 and 2.8-fold) at 3 hpt and 24 hpt after which the expression level returned to baseline (Fig. 5D).

**Figure 5.**
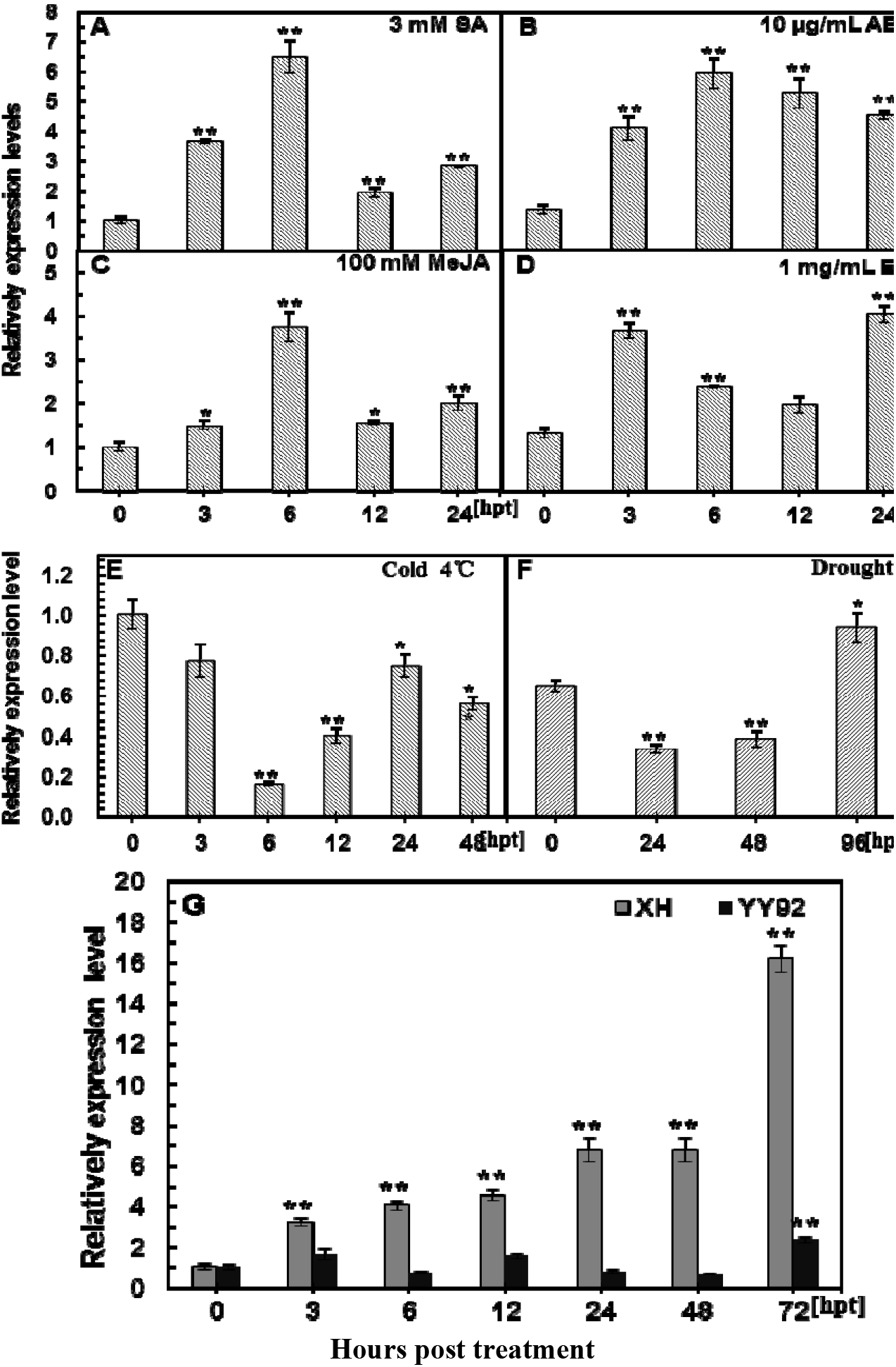
qRT-PCR analysis of *AhRLK1* transcripts in the peanut cultivar Minhua 6 under bio/abiotic treatments. Relative levels of *AhRLK1* expression in peanut leaves at different time points after treatment with (A) Salicylic acid (SA, 3 mM), (B) Abscisic acid (ABA, 10 µg/mL), (C) Ethylene (ET, 1 mg/mL), and (D) Methyl jasmonate (MeJA, 100 mM). *AhRLK1* expression performed at various hourly intervals after treatment with (E) low temperature (4 °C) and (F) drought in peanut plants at the eight-leaf stage. (G) *AhRLK1* was more up regulated in the susceptible than in the resistant variety with continuous increase as time elapsed after inoculation with *R. solanacearum*. The relative expression level of *AhRLK1* in peanut plants at various time points was compared with that in mock or control plants, which was set to 1. Asterisks indicate a significant difference (Student–Newman–Keuls test; *P < 0.05 or **P < 0.01). Error bars indicate the standard

The expression of *AhRLK1* under low temperature and drought was also examined in eight-leaf Minhua 6 seedlings (Fig. 5E and 5F). The time course of *AhRLK1* expression showed a trough in response at 6 and at 24–48 hpt upon abiotic stresses of low temperature and drought, respectively. Specifically, the transcript levels of *AhRLK1* under low temperature decreased at 3 and 6 hpt and then were up regulated between 12 and 48 hpt; the highest level of this transcript (2.5-fold) was observed at 48 hpt; however, the gene was then completely down regulated (Fig. 5E). Under drought treatment, compared with the control, the *AhRLK1* transcript level was down regulated by two-fold at 24 hpt but was up regulated at 48 and 96 hpt, with a 3.3-fold induction at 96 hpt (Fig. 5F). The transcript levels of *AhRLK1* were respectively determined by qPCR at different time points after *R. solanacearum* challenge to resistant (YY92) and susceptible (XH) peanut cultivars. In YY92, *AhRLK1* transcripts did not change within 48 h after inoculation with a highly virulent *R. solanacearum* strain. By contrast, the expression level of *AhRLK1* in XH gradually increased up to 16-fold at 96 hours post-inoculation (hpi) (Fig. 5G). This obvious transcriptional response suggested that *AhRLK1* participates in resistance to *R. solanacearum* in peanut.

### Transient overexpression of AhRLK1 in N. benthamiana leaves induced a hypersensitive response

To verify whether *AhRLK1* overexpression caused hypersensitive response (HR) cell death, 35S::*AhRLK1* was further transformed into *Agrobacterium* GV3101 and transiently expressed in *N. benthamiana* leaves by infiltration. At 48 h after infiltration, the transient overexpression of *AhRLK1* in *N. benthamiana* leaves induced an intensive HR that mimicked cell death, whereas no visible HR cell death was found in the plants infiltrated with GV3101 harbouring the empty vector 35S::00. Electrolyte leakage measurement and dark trypan blue staining showed that *AhRLK1* overexpression could trigger an HR in *N. benthamiana* leaves (Fig. 6A and 6B). DAB staining revealed high H_2_O_2_ accumulation in *N. benthamiana* leaves after *AhRLK1* overexpression (Fig. 6B). Therefore, the transient overexpression of *AhRLK1* in tobacco leaves likely induced HR and H_2_O_2_ generation as it would in response to stress.

**Figure 6.**
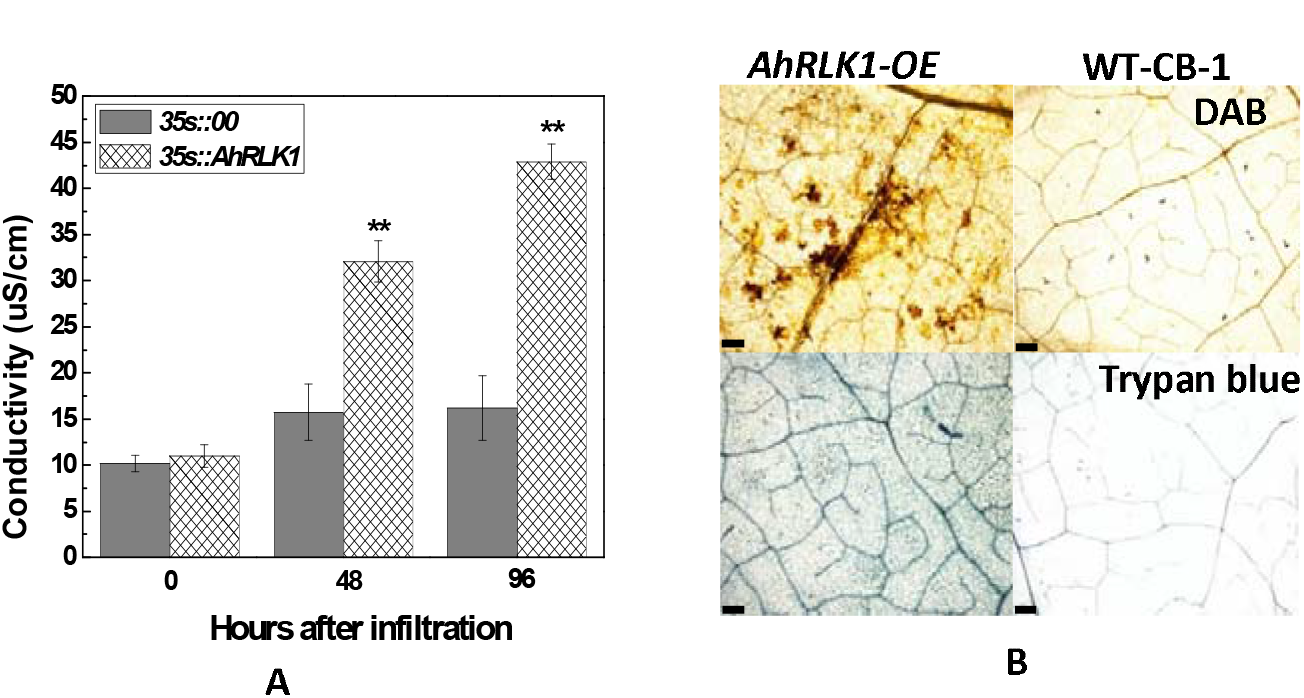
Effect of transient *AhRLK1* overexpression on immunity induction in *N. benthamiana*. (A) Electrolyte leakage of *N. benthamiana* leaves infiltrated with the *Agrobacterium* strain GV3101 containing *35S::AhRLK1* and *35S::00*. (B) Trypan blue and DAB staining of cell death and H_2_O_2_ generation, respectively, in *N. benthamiana* leaves 48 h after *AhRLK1*–*Agrobacterium* infiltration. Bars = 0.1 mm. Error bars indicate the standard error. Letters mark statistically significant differences between the wild-type and *35S::AhRLK1* tobacco by the Student–Newman–Keuls test (*P<0.05 or **P<0.01). Error bars indicate the standard error.

### Overexpression of AhRLK1 in tobacco increased resistance to R. solanacearum

To evaluate whether *AhRLK1* is involved in resistance to *R. solanacearum*, the conventional tobacco cultivar CB-1, with medium susceptibility to bacterial wilt, was transformed with *AhRLK1* driven by the *CaMV35S* promoter via an *Agrobacterium*-mediated method. The vector frame is shown in Fig. 7A. Transgenic T0 and T1 tobacco plants were generated and examined for the role of tobacco–*R. solanacearum* interaction. Compared with the wild-type cv. CB-1, the T1 transgenic generation plants of *AhRLK1-OE* showed significantly increased resistance to bacterial wilt at 40 days post-inoculation (dpi) with *R. solanacearum*. Most control plants died, with only 4 of 65 wild-type plants surviving after *R. solanacearum* inoculation. However, most transgenic plants showed high resistance to bacterial wilt, and the death rate was greatly reduced (Fig. S4). Three T2 transgenically pure lines were obtained and evaluated by inoculation with the pathogen (*AhRLK1-OE*, Fig. 7B). All of the tested transgenic lines exhibited increased disease resistance in response to *R. solanacearum* inoculation. Obvious wilting symptoms were observed on the leaves of wild-type plants at 7 dpi; whereas only slight wilting symptoms were exhibited on the *AhRLK1-OE* leaves (Fig. 7C and 7D). Severe wilting symptoms were observed in the wild-type plants at 15 dpi but not in the *AhRLK1-OE* transgenic lines. *AhRLK1* resistance was further evaluated in a hypersusceptible cultivar Honghuadajinjuan and six transgenic T_2_ homozygous lines, which were inoculated and compared with the wild type. All lines showed increased resistance to *R. solanacearum* (Fig. 7E and S5). Line 1 displayed the highest resistance with a low infection index (21.84 %) and death rate (6.80 %) at 21 dpi; however, the wild type showed serious wilting with a 95.64 % index and death rate of 86.05 % at 21 dpi (Table 1 and Table S2). This result indicated that *AhRLK1* overexpression greatly increased disease resistance against *R. solanacearum* in tobacco.

**Figure 7.**
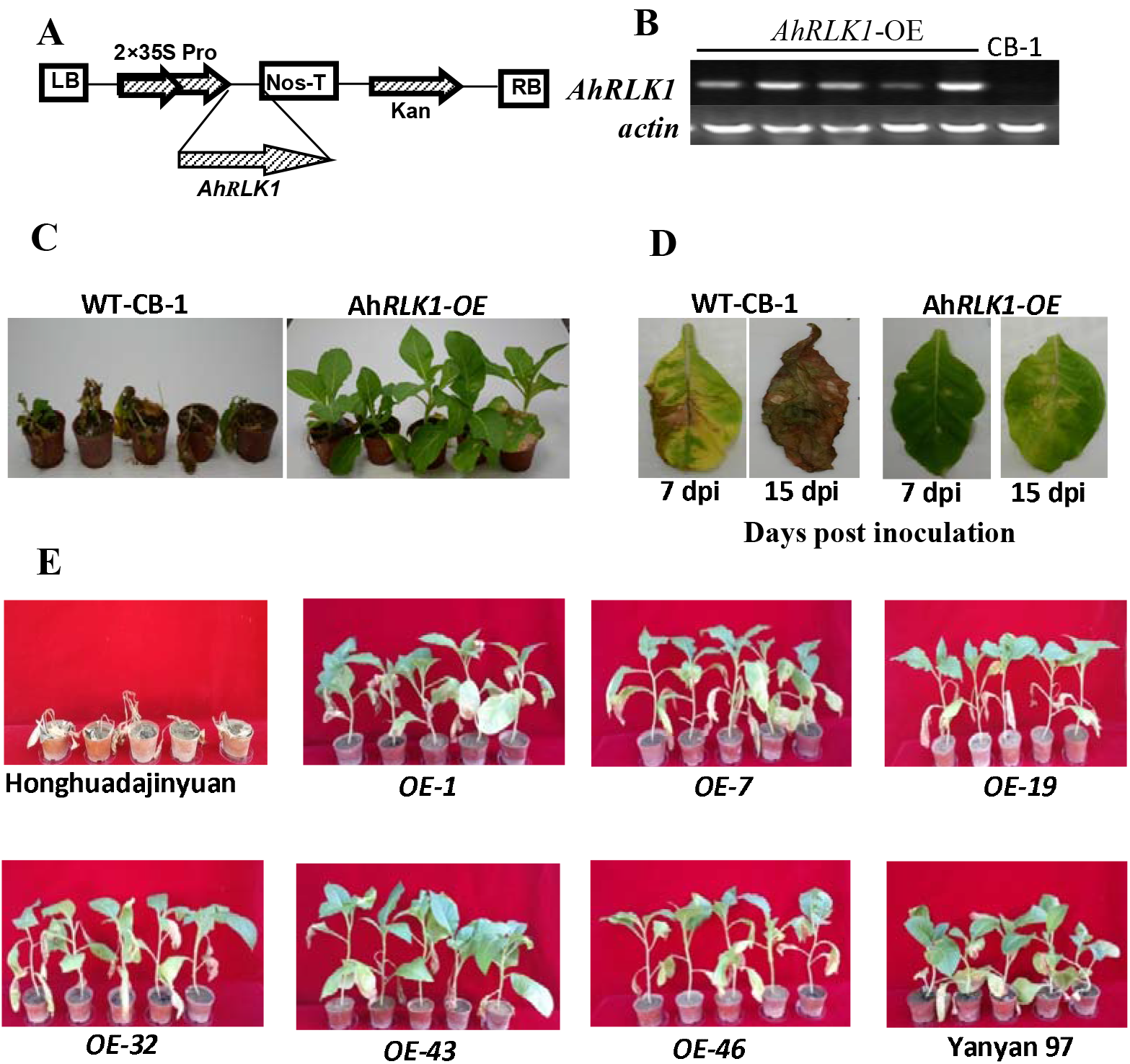
Overexpression of *AhRLK1* increased resistance to *Ralstonia solanacearum* in transgenic tobacco. (A) Schematic of the pBI121-*AhRLK1* construct. LB and RB, the left and right borders of the T-DNA; 2×35SPro, two cauliflower mosaic virus 35S promoters; Nos-T, nos-terminator; Kan^r^, kanamycin resistance. (B) RT-PCR analysis of *AhRLK1* expression in transgenic and wild-type tobacco plants; the expression level of *NtActin* was visualized as the endogenous control. (C) Third leaves of 8-week-old wild-type tobacco (CB-1，a medium susceptible cultivar) and *AhRLK1-OE* transgenic plants inoculated with a 10 µL suspension of 10^8^ cfu/mL of a highly virulent *R. solanacearum* strain. Photos were obtained at 15 days postinoculation (dpi). (D) Disease symptoms of detached leaves of wild-type and *AhRLK1-OE* transgenic plants after inoculation with *R. solanacearum*. Transgenic leaves showed immune resistance or the highly resistant phenotype. Photos were obtained at 7 and 15 dpi. (E) Overexpression of *AhRLK1* made hyper-susceptible tobacco show significantly enhanced resistance to *Ralstonia solanacearum*. Honghuadajinyuan is the hyper-susceptible tobacco cultivar as transgenic host control; Yanyan 97 is a hyper-resistent tobacco cultivar as resistant control. OE-1, OE-7, OE-19, OE-32, OE-43, OE-46 were different transgenic lines. Photos were obtained at 15 days post-inoculation (dpi) of plants after inoculation with *R. solanacearum*. All six transgenic lines showed higher resistant phenotype compared with

### Specific marker genes were upregulated in AhRLK1-transgenic tobacco in response to R. solanacearum

To confirm the role of *AhRLK1* and elucidate its possible molecular mechanism in plant disease resistance, we examined the transcriptional responses of defense-related genes, HR-responsive genes, and marker genes for SA, JA, and ET responses in *AhRLK1-OE* transgenic tobacco and wild-type CB-1 plants during *R. solanacearum* infection (Fig. 8). As shown in Fig. 8A-D, the transcripts of HR-responsive genes *NtH1N1, NtHSR201*, and *NtHSR515* were significantly up regulated in the transgenic plants (P < 0.01 or P < 0.05) but changed much less in wild-type CB-1 to different extents at 48 h after inoculation with *R. solanacearum*. By contrast, *NtHSR203* showed down regulation in response to strain infection in both transgenic and wild-type plants (Fig. 8A). The expression levels of the SA-responsive genes *NtPR2, NtPR3* and *NtCHN50* increased in the *AhRLK1*-OE-1 plants by 36, 550 and 9-fold, respectively, and were much higher than those of CB-1 in response to the pathogen. However, *NtRP4* showed less significant up regulation in the transgenic lines in comparison with the controls (Fig. 8B). The JA-responsive genes *NtLOX1* and *NtPR1b* were both up regulated in CB-1, but the up regulation was more significant in transgenic plants in response to the pathogen, with transcripts 36- and 46-fold higher than those in non-inoculation controls. Additionally, *NtDEF1* was significantly up regulated by more than 9-fold, compared with the wild type after inoculation with the pathogen (Fig. 8C). The transcript levels of the ET-responsive genes *NtEFE26* and *NtACS6* also increased significantly at 48 h after *R. solanacearum* infection in transgenic plants, whereas in wild-type plants, the up regulation of *NtEFE26* was less, and the expression of *NtACS6* was reduced (Fig. 8D). Clearly, the expression of most pathogen-inducible genes associated with HR and hormone defense signalling increased under *AhRLK1* overexpression. However, the expression of a few genes was reduced, which was also consistent with the increase in resistance to *R. solanacearum*.

**Fig.8.**
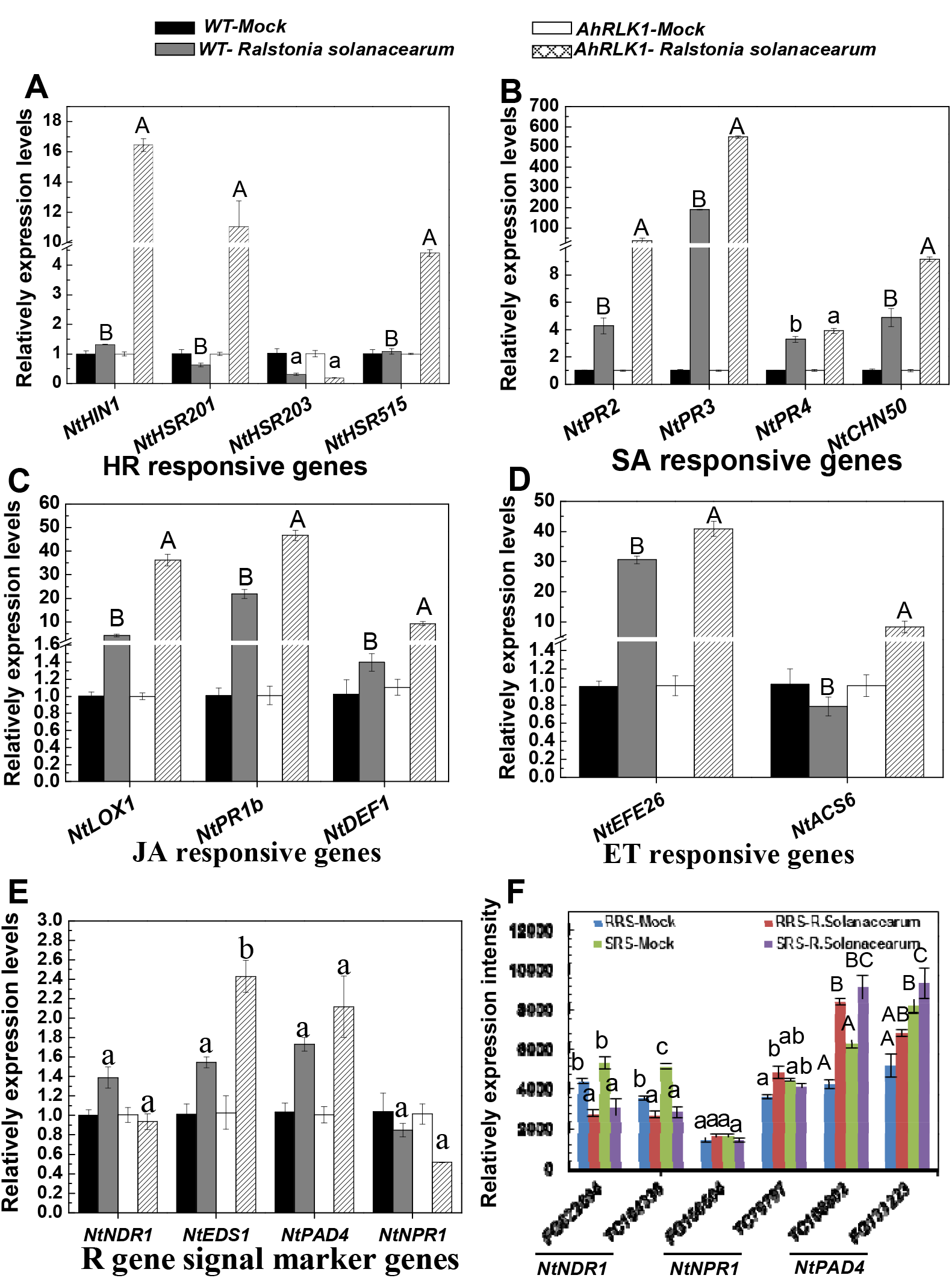
Transcript levels of the defence marker genes in transgenic or nontransgenic tobaccos and resistant and susceptible varieties after inoculation with *R. solanacearum*. A-E. The transcript levels of some defence marker genes of the *35S::AhRLK1* transgenic tobacco plants and the wild-type CB-1 by qRT-PCR. The *NtHIN1, NtHSR201, NtHSR203*, and *NtHSR515* in HR signal (A)*, NtPR2, NtCHN50, NtPR3*, and *NtPR4* in SA signal (B), *NtLOX1, NtPR-1b*, and *NtDEF1* in JA signal (C), and *NtEFE26* and *NtAsc6* in ET signal (D) pathways and *NtNDR1, NtEDS1, NtPAD4* and *NtNPR1* in R-gene resistant signal pathway (E) were determined by qRT-PCR. Transcript levels were normalized using *NtEF1*. The transcript levels of non-inoculated plants were used as the controls and assigned the value of 1. AhRLK-*R. solanacearum* and WT-*R. solanacearum* were transgenic or wild-type plants with inoculation of pathogen, respectively; AhRLK-Mock and WT-Mock were transgenic or wild-type without inoculation, respectively. F. In silico analysis of marker genes expression in R gene signal with/without inoculation of pathogen in the resistant Yueyou 97 and susceptible Honghuadajinyuan. FG622694 and TC104336 are two NDR1-like genes; FG156504 and TC79797 are NPR1/NIM1-like genes; TC108802 and FG133223 are PAD4 genes. RRS-*R. solanacearum* indicates hyper-resistant tobacco variety Yanyan 97 under inoculation; RRS-*Mock* indicates hyper-resistant variety Yanyan 97 without inoculation. SRS-*R. solanacearum* indicates hyper-susceptible variety Honghuadajinyuan with inoculation; SRS-*Mock*, The letters mark statistically significant differences between the wild type and 35S::AhRLK1 tobacco plants by the Student–Newman–Keuls test (lowercase difference mark, P < 0.05; uppercase difference mark, P < 0.01).

To further characterize the increased resistance provided by *AhRLK1* to *R. solanacearum* in transgenic tobacco, we examined several marker genes in R-gene signalling (Fig. 8E). The transcripts of *NtEDS1* and *NtPAD4* were up regulated significantly in *AhRLK1-OE* transgenic plants, with levels much higher than those in wild-type controls after inoculation of the pathogen. However, the transcripts of *NtNDR1* and *NtNPR1* declined significantly to levels lower than those in wild-type controls. We also investigated these marker genes in the resistant cultivar Yanyan97 and the susceptible cultivar Honghuadajinyuan in response to *R. solanacearum* infection with chip hybridization using non-amplified double strains of cDNA (Fig. 8E). The two members of *NDR1* genes both showed down regulation in resistant and susceptible varieties, whereas the transcripts of two *PAD4* genes increased significantly in response. Nevertheless, for the *NPR1*-like genes, down regulation occurred in the susceptible variety but up regulation in the resistant one. Therefore, the overexpression of *AhRLK1* in transgenic tobacco contributed to *R. solanacearum* resistance by involving in a series of signalling pathways, in addition to employing the *EDS1* pathway in the R-gene signalling, whereas resistance in the wild type was realized using *EDS1* and *NPR1* pathways.

## Discussion

### AhRLK1 characterized as CLAVATA1 participates in defense response to R. solanacearum

The *AhRLK1* identified from peanut by microarray hybridization as an up regulated responsive factor to *R. solanacearum* challenge was a typical LRR-RLK family gene (Torii *et al*. 1996). The full length CDS of this gene was isolated by RACE and contained 12 conserved LRRs and a kinase domain. Phylogenetic analysis showed it was a CLAVATA1-like protein, which are responsible for shoot and root meristem development, among other functions (Clark *et al*., 1993, 1997; Williams and De Smet, 2013). Microarray anal ysis indicated three genes in the *AhRLK1* family were all expressed most strongly in stem and roots, but only traces were found in the embryo or in the pericarp, suggesting their roles in root and stem development (Alvarez *et al*., 2013; Williams and De Smet, 2013; Hazak and Hardtke, 2016). It also showed high similarity with several known functional LRR-RLK genes, such as *FLS2* identified from flg22-sensitive *Arabidopsis* mutants, which shows receptor activity that can induce pathogen response (Gómez-Gómez and Boller, 2000; Gómez-Gómez *et al*., 2001), and *OSXa21*, a resistance gene of rice, which specifies the gene-for-gene resistance of rice against *Xanthomonas oryzae* (Song *et al*., 1995; Wang *et al*., 1996). The *AtERECTA* is another *Arabidopsis* LRR-RLK gene resistant to *R. solanacearum* (Godiard, Laurence and Sauviac, Laurent and Torii, Keiko U and Grenon, Olivier and Mangin, Brigitte and Grimsley, Nigel H and Marco, 2003). Real-time PCR showed it was up regulated with time in response to *R. solanacearum* inoculation in Xinhuixiaoli but remained almost unchanged in Yueyou92. Therefore, AhRLK1 might not only function similarly to the *CLAVATA1* associated with shoot meristem determination but also participate in the defense response to *RS* infection. In a recent study, *Atclv1*, a mutant of *CLAVATA1*, increased the resistance to *RS* in *Arabidopsis* (Hanemian *et al*., 2016), consistent with *AhRLK1* as an *RS* defense responsive factor.

### AhRLK1 is widely associated in defense responses to biotic/abiotic stresses

The LRR-RLK gene family participates widely in the regulation of plant growth and development and also in the resistance to pathogens and environmental stresses (Clark *et al*., 1993; Godiard, Laurence and Sauviac, Laurent and Torii, Keiko U and Grenon, Olivier and Mangin, Brigitte and Grimsley, Nigel H and Marco, 2003; Sun *et al*., 2004; Wu *et al*., 2009; Xu *et al*., 2009; Hanemian *et al*., 2016). Both *AtCLV1* and *AtCLV2* in *Arabidopsis* are involved in meristem identity, and in their mutants, *clv1* and *clv2*, the resistance to bacteria pathogens increases. Their resistance did not require the CLV signalling modules involved in meristem homeostasis and was not conditioned by defense-related hormones such as salicylic acid, ethylene, and JA (Hanemian *et al*., 2016). In peanut, we found that *AhRLK1* responded differentially to *R. solanacearum* inoculation in resistance and susceptible varieties (Fig. 5G). The transcript of *AhRLK1* was up regulated by the treatment of hormones such as SA, ABA, JA and ET, although with slightly different expression patterns. However, the response patterns of transcripts to cold and drought stress were completely different (Fig. 5A-F). Clearly, the expression of peanut *AhRLK1* was differentially affected with exposure to various hormones and environmental stresses. Both *Arabidopsis* and soybean *CLV1* function as receptor subunits in the CLAVATA2/CORYNE (CRN) heterodimer complex; although, receptor-like protein kinase 2 is required for perception of CLEs, secreted from the nematodes *Heterodera schachtii* and *Heterodera glycines* (Guo *et al*., 2015). The expression of *AhRLK2* is induced at the feeding sites of these nematodes on roots. Mutants of *CLV1* show increased resistance to the nematodes in soybean. However, *AhRLK1*, the ortholog of *Arabidopsis AtCLV1*, is widely involved in defense response to biotic stress and in shoot and root meristem homeostasis.

### AhRLK1 confers resistance to bacterial wilt in transgenic tobacco

*AtRRS1-R*, the first characterized resistant gene to *R. solanacearum*, was a specific TIR-NBS-LRR gene containing a WRKY domain at the C-terminal (Deslandes *et al*., 2002). Transgenic *Arabidopsis* overexpressing the recessive *RRS1-R* conferred dominant resistance to *R. solanacearum* GMI1000 (Deslandes *et al*. 2003). The *RPS4* was later identified associated with *RRS1-R* for the resistance to bacterial wilt and also to other two diseases (Gassmann *et al*., 1999; Narusaka *et al*., 2009; Sohn *et al*., 2014). Additionally, a QTL, named *ERECTA*, was isolated as an LRR-RLK gene that showed resistance to bacterial wilt and regulation in the development of aerial organs (Godiard *et al*. 2003).

In our study, transient expression of AhRLK1::GFP fusion protein in *N. benthemiana* showed AhRLK1 localized at the membrane and cytoplasm (Fig. 3) at which it functions. With overexpression of *AhRLK1* in a medium susceptible tobacco cultivar, CB-1, the resistance to bacterial wilt increased significantly. Furthermore, 6 different transgenic T_2_ homozygous lines derived from the hypersusceptible tobacco cultivar Honghuadajinyuan and carrying an overexpression *cassette of AhRLK1* also showed significantly increased but diverse levels of resistance to *R. solanacearum* (Fig. 7E, S5, Table 1). These lines demonstrated that *AhRLK1* could confer resistance to bacterial wilt in a heterogeneous crop. Transient overexpression of *AhRLK1* in *N. benthemiana* suggested a hypersensitive response was induced, based on trypan blue staining and DAB accumulation and also the production of H_2_O_2_, which indicated *AhRLK1* could result in the cell death caused by hypersensitive response. Thus, the implication was that *AhRLK1* might employ an ROS pathway for its resistance.

However, the mechanism of AhRLK1 is apparently different from that of *Atclv1, Atclv2* and *Atcrn* mutants, which are null alleles and different genes and all show increased resistance via a decrease in miR169 accumulation (Diévart *et al*. 2003; Hanemian *et al*. 2016). Wild-type genotypes including *AtCLV1, AhCLV2* and *CRN* demonstrate susceptible phenotypes (Hanemian *et al*. 2016). By contrast, AhRLK1 is a functional gene. *AhRLK1* expression changed following treatments of hormones such as ABA, Eth, and SA, which suggested that AhRLK might confer resistance to *R. solanacearum* through other mechanisms different from those of *AtCLV1* and *AtCLV2*, with their mutants that increase resistance via a defect in miR169 accumulation (Hanemian *et al*., 2016). Therefore, our report is the first that a peanut RLK is involved in resistance to *R. solanacearum* and confers resistance in a heterologous crop.

### AhRLK1 resistance is associated with the R gene and defense signalling in transgenic tobacco

A complex network of many defense signalling pathways are involved in plant-pathogen interactions, each of which is associated with some marker genes in their mediated disease resistance reaction (Divi *et al*. 2010; Nahar *et al*. 2012; Yang *et al*. 2013; Vos *et al*. 2015;). In the comparison between the *AhRLK1-OE* and wild-type tobacco variants in association with *R. solanacearum* based on real-time PCR, marker genes *NtHIN1, HSR201*, and *HSR515* in *HR* signalling (Sohn *et al*., 2007) were significantly activated in transgenic lines under inoculation of the pathogen (Fig. 8A). This result was consistent with the phenotype of transient overexpression of *AhRLK1* in *N. benthemiana*, which led to HR and cell death (Fig. 6A, B), thereby indicating that the resistance employed HR signalling. Some PR genes, such as *NtPR2, NtPR3*, and *NtCHN50* in SA signalling, were highly induced in overexpressed lines of *AhRLK1* (Dong, 1998; Glazebrook, 2005), suggesting that SA signalling was also associated with the AhRLK1 resistance. The ET signalling marker genes *NtACS6* and *NtEFE26* and the JA signalling genes *NtPR1b, NtDEF1* and *NtLOX1* were all up regulated in overexpressed lines of *AhRLK1* (Fig. 8). This result was consistent with those observed in peanut in which *AhRLK1* was up regulated by the exogenous applications of SA, ET, JA, and ABA. Based on these lines of evidence, the interplay of different hormones signals is implicated in the increased resistance of transgenic tobacco with peanut *AhRLK1*. In rice, XA21 is a receptor-like kinase that confers resistance against most strains of Xoo (Song *et al*., 1995). SA is required for *XA21*-mediated full resistance to *Xoo*, and the resistance to Xoo decreases but is not completely abolished in *Xa21/NahG* plants (Lee *et al*., 2009). However, *Atclv1, Atclv2* and *crn1* mutants of *AtCLV1, AtCLV2* and *CRN1*, respectively, all showed increased resistance to bacterial wilt, which apparently did not require hormone signalling, such as that from ABA, ET, JA and SA. Therefore, the resistance of peanut mediated by *AhRLK1* could be different from that of the orthologs *Atclv1* and the *Atclv2 and Atcrn1*. Because the gene is for meristem determination, the mechanism by which *AhRLK1* employs multiple hormones in fine-tuning immune responses in peanut requires further study.

*NDR1* and *EDS1* are important regulators for R-gene-mediated resistant signalling in plants (Day *et al*., 2006; Bhattacharjee *et al*., 2011; Lu *et al*., 2013). Usually, *NDR1* is involved in the resistance mediated by CC-NBS-LRR-type of R genes, and *EDS1* and *PAD4* are implicated in Tir-NBS-LRR resistant signalling (Aarts *et al*., 1998; Wang *et al*., 2014). However, *RRS1-R*, a Tir-NBS-LRR gene in *Arabidopsis*, and *AhRRS5*, an NBS-LRR gene in peanut, require *NDR1* for their resistance (Deslandes *et al*., 2002; Zhang *et al*., 2017). In this study, overexpression of *AhRLK1* in transgenic tobacco reduced *NDR1* expression, but more significantly, the transcripts of *EDS1* and *PAD4* were up regulated in response to *R. solanacearum*, compared with those in the wild type. However, it down regulated the transcripts of *NPR1* in the transgenic plants responding to the pathogen (Fig. 8E). NPR1 is a key regulator of SAR and is essential for SA signal transduction to activate PR gene expression associated with R-gene resistance (Pieterse and Van Loon, 2004; Sandhu *et al*., 2009; Xia *et al*., 2013). Thus, the results indicated that *AhRLK1* was associated with the *EDS1* pathway in the R-gene signal for the resistance to the pathogen in transgenic tobacco, although NPR1 was not required for this resistance.

For comparison, in silico hybridization with double strains of cDNA showed that the expression of two *NDR1* genes declined in hyperresistant non-transgenic Yanyan97 in response to the pathogen. By contrast, *PAD4* genes in the *EDS1* pathway were up regulated in response to the pathogen, which was a phenomenon consistent with the transgenic tobacco overexpressing *AhRLK1*. However, the expression of the two *NPR1* decreased in the hypersusceptible cultivar but was up regulated in the hyperresistant variety in response to the pathogen. This result is consistent with the report of *NPR1*-mediated resistance to viral and bacterial pathogens and that repressing *NPR1* transcripts increases the susceptibility of plants to pathogens (Xiao and Chye, 2011; Li *et al*., 2012). By contrast, in this study, with high resistance conferred to *R. solanacearum* by *AhRLK1*, the expression of *NPR1* was down regulated. Thus, we further suggest that *AhRLK1* participated in pathogen resistance by employing the R-gene pathway in association with *NtEDS1* but independent of *NtNPR1*.

## Supplementary data

Additional supporting information is in the online version of this article:

**Supplementary Fig. S1.** Cloning of *AhRLK1* from peanut. Electrophoresis photos represent 5’ RACE, 3’ RACE and full-length cDNA PCR product of *AhRLK1*.

**Supplementary Fig. S2.** Multiple sequence alignment with known functional LRR receptor kinase proteins.

**Supplementary Fig. S3.** Phylogenetic tree constructed using AhRLK1 and 180 different subfamily LRR RLKs of *Arabidopsis*.

**Supplementary Fig. S4.** Phenotype of *AhRLK1-OE* transgenic T1 lines and non-transgenic control plants in tobacco cultivar CB-1 after inoculation with *R. solanacearum* for 40 days.

**Supplementary Fig. S5.** Resistance phenotype of T_2_ *AhRLK1-* OE transgenic homozygous lines and the control plants.

**Supplementary Data S1.** Sequences of *AhRLK1* full-length cDNA, genomic DNA, and protein.

**Supplementary Data S2.** Amino acid sequences of five homolog LRR-RLKs.

**Supplementary Data S3.** Thirty-five known functional *Arabdopsis* LRR-RLK proteins used for phylogenetic analysis.

**Supplementary Data S4.** In silico study of expression characteristics of three members in the *AhRLK1* family in peanut.

**Supplementary Table S1.** Primary primers for PCR used in this study.

**Supplementary Table S2.** Detailed data of disease indexes and death ratios of different OE lines and the wild type after inoculation with *Ralstonia solanacearum*.

## Acknowledgements

This work was supported by The educational and scientific research program for young and middle-aged instructor of Fujian province (JAT170165), The National Science Foundation of P. R. China (U1705233; 31701463).

